# MESH-FREE HIGH-RESOLUTION SIMULATION OF CEREBROCORTICAL OXYGEN SUPPLY WITH FAST FOURIER PRECONDITIONING

**DOI:** 10.1101/2023.01.09.523320

**Authors:** Thomas Ventimiglia, Andreas A Linninger

## Abstract

Oxygen transfer from blood vessels to cortical brain tissue is representative of a class of problems with mixed-domain character. Large-scale efficient computation of tissue oxygen concentration is dependent on the manner in which the tubular network of blood vessels is coupled to the tissue mesh. Models which explicitly resolve the interface between the tissue and vasculature with a contiguous mesh are prohibitively expensive for very dense cerebral microvasculature. We propose a *mixed-domain mesh-free* technique whereby a vascular anatomical network (VAN) represented as a thin directed graph serves for convection of blood oxygen, and the surrounding extravascular tissue is represented as a Cartesian grid of 3D voxels throughout which oxygen is transported by diffusion. We split the network and tissue meshes by the Schur complement method of domain decomposition to obtain a reduced set of system equations for the tissue oxygen concentration. The use of a Cartesian grid allows the corresponding matrix equation to be solved approximately with a fast Fourier transform based Poisson solver, which serves as an effective preconditioner for Krylov subspace iteration. The performance of this method enables the steady state simulation of cortical oxygen perfusion for anatomically accurate vascular networks down to single micron resolution without the need for supercomputers.

**Practitioner Points:**

- We present a novel mixed-domain framework for efficiently modeling O_2_ extraction kinetics in the brain.
- Model equations are generated by graph-theoretic methods for mixed domains.
- Dual mesh domain decomposition with FFT preconditioning yields very fast simulation times for extremely high spatial resolution.

## 1 Introduction

Metabolic studies in normal or aging brains have motivated the construction of computational models for oxygen transfer from cerebral blood supply to brain tissue [1]. The reliability of physiological simulation depends on its ability to model blood flow and tissue interactions for realistic vascular networks at all length scales, yet the dense and tortuous nature of the capillary bed poses a massive computational challenge. For example, a cortical sample of 1×1×1 mm^3^ at 1 μm resolution requires 1 billion equations.

### Single capillary models

Classical approaches to the modeling of O_2_ perfusion focused on analytical solution of a differential equation for a single capillary with prescribed boundary conditions: the Krogh model solves for the oxygen tension in a cylindrical annulus of tissue surrounding a single straight capillary along its centerline [2]. This solution was applied to the context of capillaries in striated muscle tissue, which are approximately parallel, and cannot be readily extended to simulations of the entire capillary bed. Furthermore, this model neglects diffusion in the axial direction, assumes constant metabolic rate, and implies that oxygen saturation decreases linearly along the capillary axis [3, 4]. Later models have attempted to use a single capillary solution together with anatomically accurate statistical distributions of capillary lengths and diameters to understand O_2_ perfusion at a larger scale: in [5] and [6], the solution for a single capillary is used to analyze blocks of tissue at the scale of voxels in fMRI imaging.

### Green’s function methods

Under the assumption that oxygen transport in tissue can be modeled by diffusion in a porous medium, Secomb et. al. [7] derive the analytic solution of the steady-state diffusion equation in the vicinity of a point source (a Green’s function) and then represent the oxygen tension in the tissue surrounding a VAN (vascular anatomical network) as a superposition of Green’s functions arising from point sources distributed along the surfaces of cylindrical blood vessels and sinks throughout cuboidal tissue elements. This method resolves the interactions between vessels and tissue in any arbitrary configuration and is superior to a naïve implementation of finite-differences [8]. Despite its utility [9], the Green’s function method is severely limited by the size of the network and number of tissue elements under consideration: it requires computing the influence of each source on every other source, and thus has a complexity of at least *O*(*n*^2^) with respect to the number of finite elements.

### Frontal finite element method

Fang et. al. [10], proposed a fully coupled model of cerebral oxygen transfer based on a triangular mesh for the vasculature that was contiguously embedded in a tetrahedral volumetric mesh for the tissue space. However, contiguous computational meshes that resolve capillaries and extravascular tissue space require an excessively large number of elements [1]. Consequently, the method becomes intractable for simulating representative samples of the cerebral cortex (> 0.5 mm^3^) at high resolution, because the dense tortuous nature of the microvasculature requires excessive number of elements to resolve the blood tissue interface (bloods brain barrier).

### Dual mesh techniques

Linninger and colleagues [11, 12] introduced *dual mesh techniques* that do not require exact resolution of the capillary-tissue interface which severely limits frontal FEMs. A *thin* (i.e. reduced dimension, 1D) vascular network graph was embedded in a coarse tissue domain (three-dimensional volume, 3D) without the need for constructing a contiguously fitted mesh. Continuity of O_2_ tension and transfer fluxes at the interface ensure consistent prediction of the radial and axial O_2_ gradients despite marked length scale differences between the two domains. The treatment of length scale separation in the proposed dual mesh technique is similar to the Lagrange multiplier approach for mixed domain PDEs in 1D-3D coupled domains [13]. Massive simulations of O_2_ extraction in the human and mouse cortex [11,14] used unstructured tetrahedral meshes. Hartung et. al. [1] substantially improved computational efficiency by proposing a mesh free Cartesian technique for representing blood and extravascular tissue. Each cuboid mesh cell or *voxel* is registered as belonging to vascular lumen, vessel wall or extravascular tissue according to its position relative to the blood lumen. Vessel walls or voxels large enough to encompass entire capillaries act as sources to supply oxygen to adjacent tissue cells. The granularity of the voxel mask can be tuned to desired microscale resolution, while interfaces at larger scales are continuously captured. The spatial separation circumvents excessively fine grain mesh detail at the smallest scales thus averting computational intractability. Moreover, the Cartesian framework does not need to store grid coordinates, thus it is termed *mesh-free*.

### Ultrafast voxelized dual-mesh method with Fast Poisson Preconditioning

The current work builds upon the foundation set in [1] to further enlarge solvable problem sizes at substantially reduced computational cost. This expansion is chiefly due to the benefits associated with a structured Cartesian domain. The banded structured linear system of the discretized diffusion equations will be shown to be amenable to highly efficient preconditioning techniques using discrete sine transforms (DST). Our method enables fully coupled nonlinear simulations of oxygen transport between blood and brain at high resolution, approaching 1 μm length scale or more than one billion equations.

## 2 Models and Methods

### 2.1 Overview

Steady oxygen perfusion in Cartesian coordinates can be cast as a Poisson problem for which *fast* matrix decomposition algorithms are available [15]. *Fast* solvers exploit the fact that the eigen-decomposition of the finite volume discretization of the Poisson equation is analytically known, so instead of elimination steps, solutions can be directly evaluated using the *discrete sine transform* (*DST*), which can be executed in *O*(*n* log *n*) operations, where *n* is the number of tissue elements [16]. Direct DST solution of the diffusion problem enables extremely efficient simulation of oxygen exchange even for problems exceeding one billion tissue elements. Since the eigenvalue decomposition and Poisson solution is analytical and direct, the explosion of the condition number [17] hampering iterative algorithms at fine mesh resolutions does not occur.

The Poisson approach, however, does not apply to convective transport of blood oxygen. We therefore chose to incorporate oxygen supply to tissue by introducing source terms into the Poisson problem that are implicitly coupled to blood oxygen tension. This approach, which implicitly eliminates unknown blood states, has the same effect as domain decomposition. For mass transfer, we implemented two models: first, linear mass transfer assumes flux across the blood brain barrier to be directly proportional to the local difference of pO_2_ concentrations between blood and tissue. In the second case, bound O_2_ must first dissociate from hemoglobin into plasma before it is available to be transferred across the BBB. In this latter case, the concentration of free O_2_ is related to the concentration of bound O_2_ via the nonlinear Hill equation [18]. Linear mass transfer with the Schur complement system [19] will be solved iteratively to compute the oxygen exchange across the BBB and then to recover the vessel states by convection. We propose that transport in tissue can be effectively preconditioned by solving a nearby (in the Frobenius norm) Poisson problem with a fast matrix decomposition algorithm [20, 21]. For non-linear oxygen dissociation kinetics, we implement Newton’s method with quadratic convergence [22].

#### Ethics Statement

All protocols were approved by the Institutional Animal Care and Use Committee at University of California, San Diego.

### 2.2 Graph representation of vascular and tissue domains

#### 2.2.1 Representation of the blood vessels – vascular graph

We regard the vascular network (VAN) as being composed of several right circular cylindrical segments with prescribed diameters. The VAN thus corresponds to a weighted directed graph whose edges represent cylindrical blood vessel segments. Each node of the graph has a finite volume element composed of the conjunction of halves of all cylinders associated with edges incident to the node. Cylinder ends at the node are covered by spherical endcaps (Figure 1). In this way we associate the entire VAN with a collection of finite volume elements whose union contains the graph in its interior. Inlets and outlets (boundary nodes) are easily identified as nodes with only one edge connection. State variables associated with the network (like blood pressure or blood O_2_ concentration) are constant throughout each well-mixed control volume. Consequently, the state variables are vectors of length *n_p_*, where *n_p_* is the number of graph nodes (=control volumes). Details on vascular data acquisition [23,24,25] and image reconstruction [26] can be found elsewhere.

**Figure 1.**
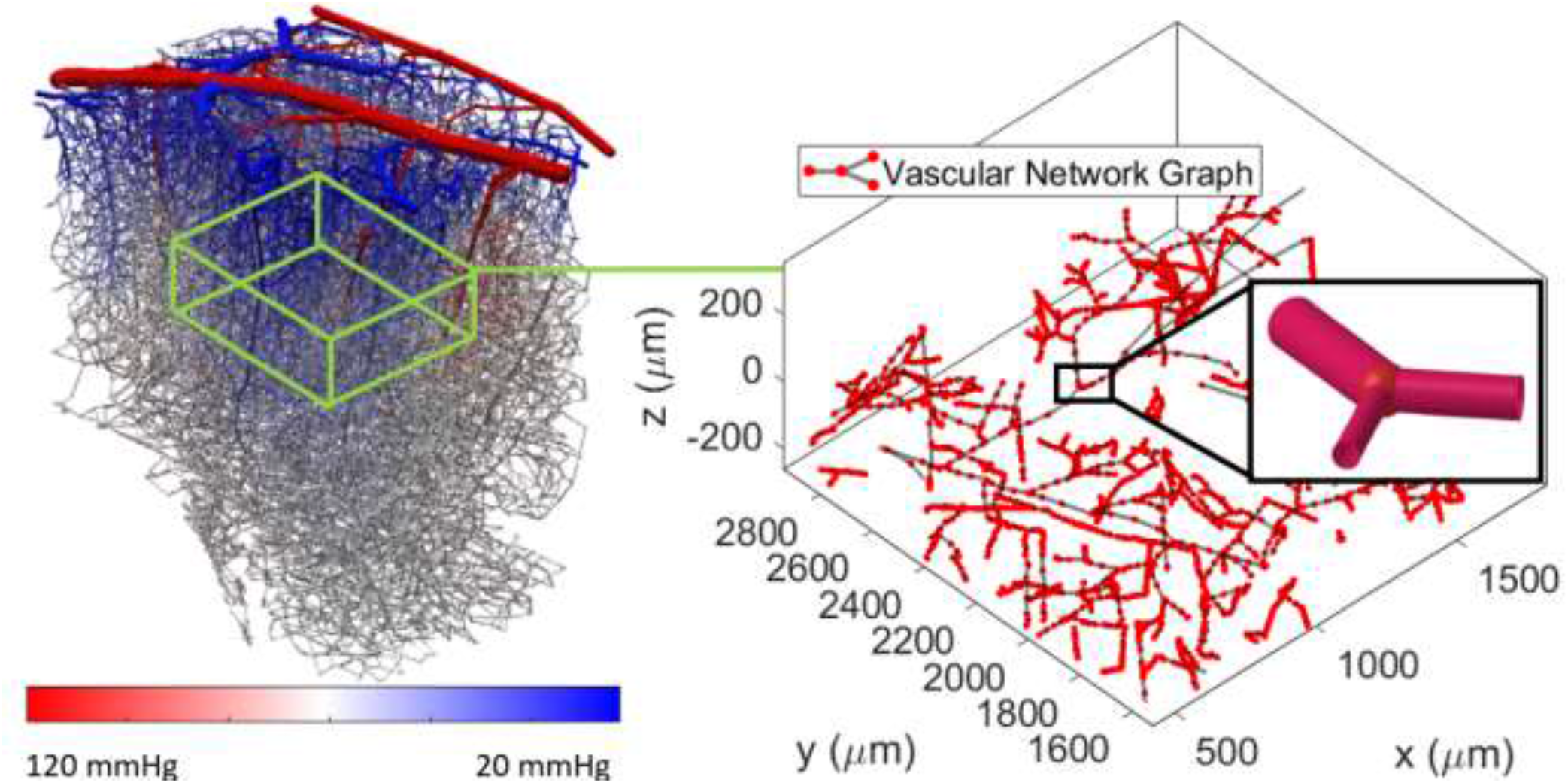
A vascular anatomical network (VAN) is represented in data as a graph composed of nodes and edges. Each edge is regarded as a right cylindrical tube with a prescribed diameter and each node is associated with a control volume formed from half cylinders with a spherical cap at the junction. Inset: the control volume associated with a bifurcation.

#### 2.2.2 Representation of the extravascular space

The tissue domain is an array of cuboid voxels occupying a 3D region of space large enough to contain the entire VAN (all nodal control volumes) in its interior. We assume that state variables defined on the tissue domain are constant throughout each voxel. Consequently, we represent the state variables as vectors of length *n_t_*, where *n_t_* is the number of tissue voxels. To be consistent with the graph representation of the vascular domain, we give a directed graph representation to the tissue domain as well by taking the nodes to be the center-points of each voxel and the edges to be directed line segments parallel to the positive *x, y*, and *z* axes connecting each node to its (at most) six neighbors through the square voxel sides. In this view, brain tissue is a three-dimensional grid graph [27].

#### 2.2.3 Voxelized mixed-domain representation

We identify the vascular network with a subset of the 3D voxel array to explicitly delineate the locations of O_2_ mass transfer in space. Following [1] we classify each voxel as belonging to either the extravascular space, the intravascular lumen, or the vessel endothelial wall (Figure 2). In this way we embed a voxelized version of the VAN in the tissue domain to obtain a mixed-domain representation of the vascular and tissue states in a common 3D space. This is to avoid explicit meshing of the vascular surface: the use of a Cartesian voxel array means that there is no need to store the coordinates of the tissue mesh as it is naturally indexed by a 3D array. The only storage requirements for the coupled domain are the VAN graph and the locations of voxels associated with the VAN, which is sparse (a trinary mask). This was referred to as a “mesh-free” method in [1]. The Cartesian mesh also enables the use of a fast Fourier transform based diffusion solver (which requires a rectangular domain) as a preconditioner for the steady-state O_2_ simulation. This is the core of our solution technique in section 2.6. See Supplemental File S1 for a detailed description of how the voxelized network is constructed.

**Figure 2.**
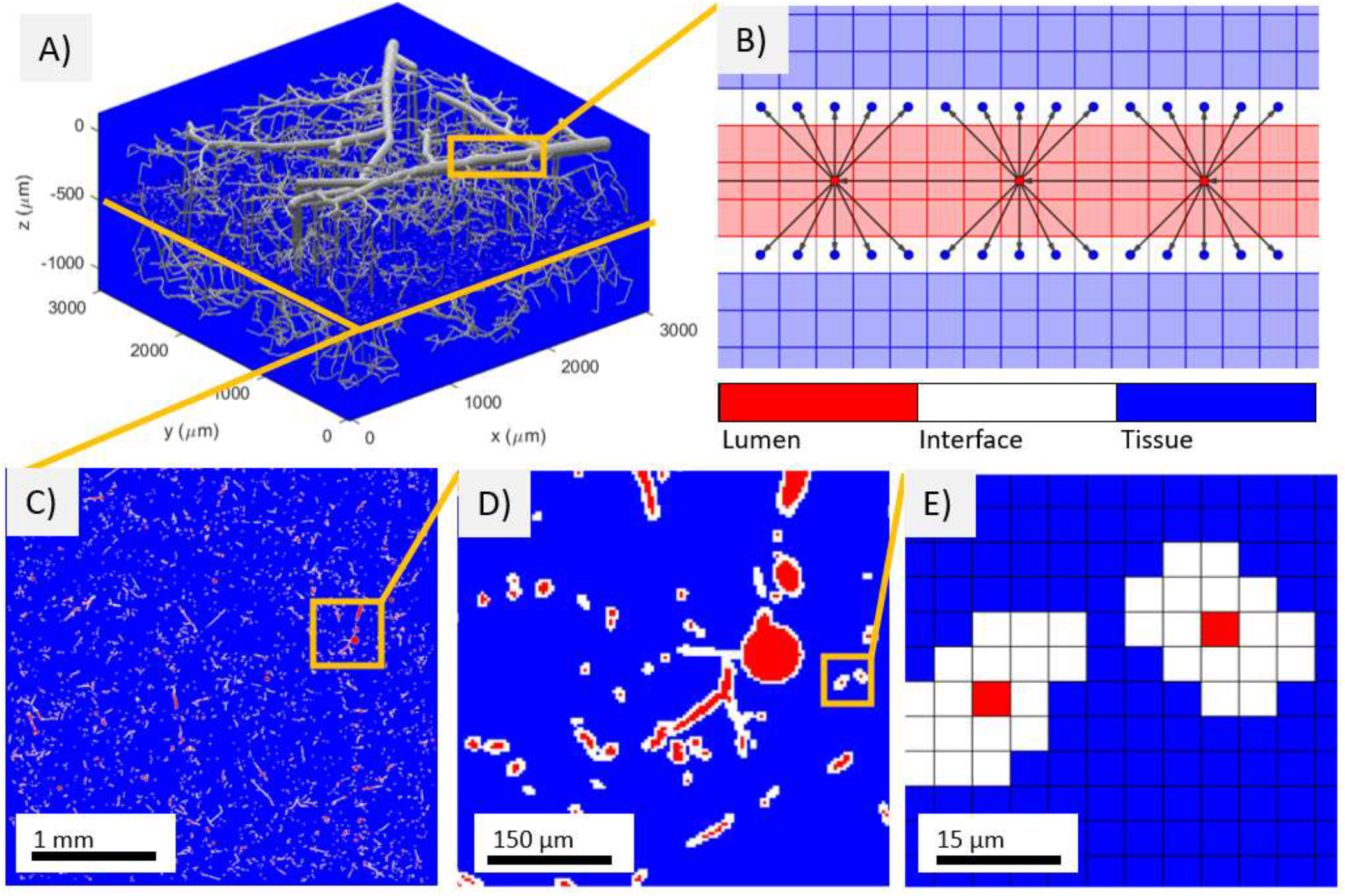
A) Voxelized network mask with smallest vessels (diameter < 12 μm) omitted for clarity. B) Schematic illustration of network coupling to tissue mesh: this is a nearest neighbor approach, where voxels comprising the vessel wall receive oxygen from the vascular network node closest to it. Alternative approaches not explored in this work are to supply each voxel on the vessel wall with a convex combination of the oxygen concentration values in nearby network nodes, or an upwinding formulation where the voxels are supplied by the nearest upstream nodes. C-E) Horizontal slice of full mask at cortical depth *z* = −500 μm at different magnifications. The constant *z* slice in panel E has cut into the small vessels at an oblique angle, causing the apparently thick vessel wall.

#### 2.2.4 Mass transfer between the vascular and tissue domains

We adjoin the graphs of the vascular and tissue domains into a single unified graph by including a directed edge (*i, j*) whenever the tissue voxel *j* ∈ {1, 2, …, *n_t_*} belongs to the set of surface (=vessel wall) voxels associated to the control volume of the VAN node *i* ∈ {1, 2, …, *n_p_*} (Figure 2, panel B). This is equivalent to adjoining to the vascular and tissue graphs the bipartite graph [28] whose distinct node sets comprise the vascular nodes and surface voxel nodes respectively and has an edge whenever a VAN node is associated with a surface voxel node. Hence for vessels with large diameter relative to the voxel side length each VAN node has multiple edges connecting it to adjacent voxel nodes, and for vessels with small diameter many node control volumes may be contained in the interior of a single voxel, so there are multiple edges connecting that voxel to the VAN nodes within it.

### 2.3 Simulation of blood flow in the vascular network

Each vessel segment is approximated as a right circular cylinder; consequently, we approximate blood flow in each edge by the Hagen-Poiseuille law for laminar fluid flow in a pipe [14].

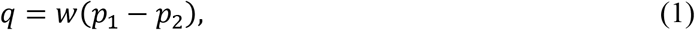

where *q* is the volumetric flow rate through the vessel, *p*_1_ – *p*_2_ is the pressure difference between the ends of the vessel (since the vessel is given by a directed edge, we may take *p*_1_ and *p*_2_ to be the pressures in the control volume of the initial and terminal node respectively), and *w* is the reciprocal of the hydraulic resistance:

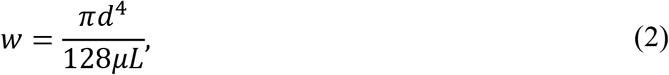

where *d* and *L* are the diameter and length of the vessel and *μ* is the (assumed constant) dynamic viscosity of blood plasma.

By introducing the *n_f_* × *n_p_* connectivity matrix *C*_1_, we can write the length *n_f_* (= number of graph edges) vector **q** of flows as the matrix equation

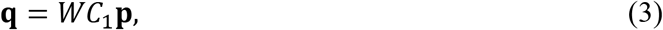

where **p** is the length *n_p_* vector of node pressures (pressure in the control volume) and *W* is the diagonal matrix of *w* values for each vessel. The matrix *C*_1_ is the transpose of the incidence matrix of the vascular graph [29]; that is, each column corresponds to a node, each row corresponds to an edge, and contains 1 in the position of the initial node and –1 in the position of the terminal node. The connectivity matrices of graphs will feature prominently in our equations since they efficiently encode the local structure of the network. See supplemental file S2 for more detail on the connectivity matrix and its use in constructing the matrix equations of this section.

At the boundary nodes we assign prescribed pressure values:

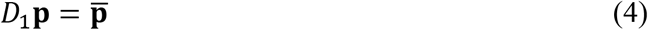

where 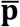 is a length *n_p_* vector with prescribed boundary values in the positions of the boundary nodes and the decision matrix *D*_1_ is an *n_p_* × *n_p_* identity matrix with ones replaced by zeros everywhere except in the positions of the boundary nodes.

We require conservation of mass in each nodal control volume. This means that for each interior node (i.e., those nodes with indegree and outdegree ≥ 1), the inflow must be balanced by the outflow:

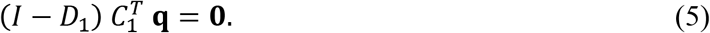

Here, *I* is an *n_p_ × n_p_* identity matrix. Prepending the equation (5) with *I* – *D*_1_ has the effect of enforcing flow balance on the interior nodes.

Combining (5) and (3) we get the following equation for the pressures at the interior nodes:

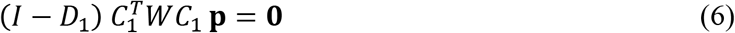

Combining (6) and (4), we get the full equation for pressures at all nodes of the vascular network:

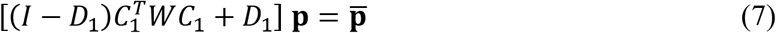

The value of the decision matrix *D*_1_ in the preceding equations is in allowing us to enforce explicit boundary conditions in a fully declarative manner, without the need for hardcoded constraints in the simulation code. While we only look at Dirichlet boundary conditions in this paper, Von Neumann or Robin boundary conditions can be enforced in a similarly declarative way, see [30] for more details on decision matrices.

By setting high pressure values at boundary nodes associated with vascular inlets (arteries) and low pressures at outlets (veins), the equation (7) solves uniquely (Figure 3, top row). For all our physiological networks, we encountered no numerical problems solving this equation directly. In every case the equation is solved in less than one minute. It is possible that numerical difficulties might arise in large vascular networks with large differences in edge resistances. For such cases, we note that the matrix on the left side of (7) is known as a *graph Laplacian* for the vascular network graph with prescribed boundary values and edge weights given by the *w* values. There is a wealth of literature on the topic of solving equations with graph Laplacians, like for example in [31] and [32].

**Figure 3.**
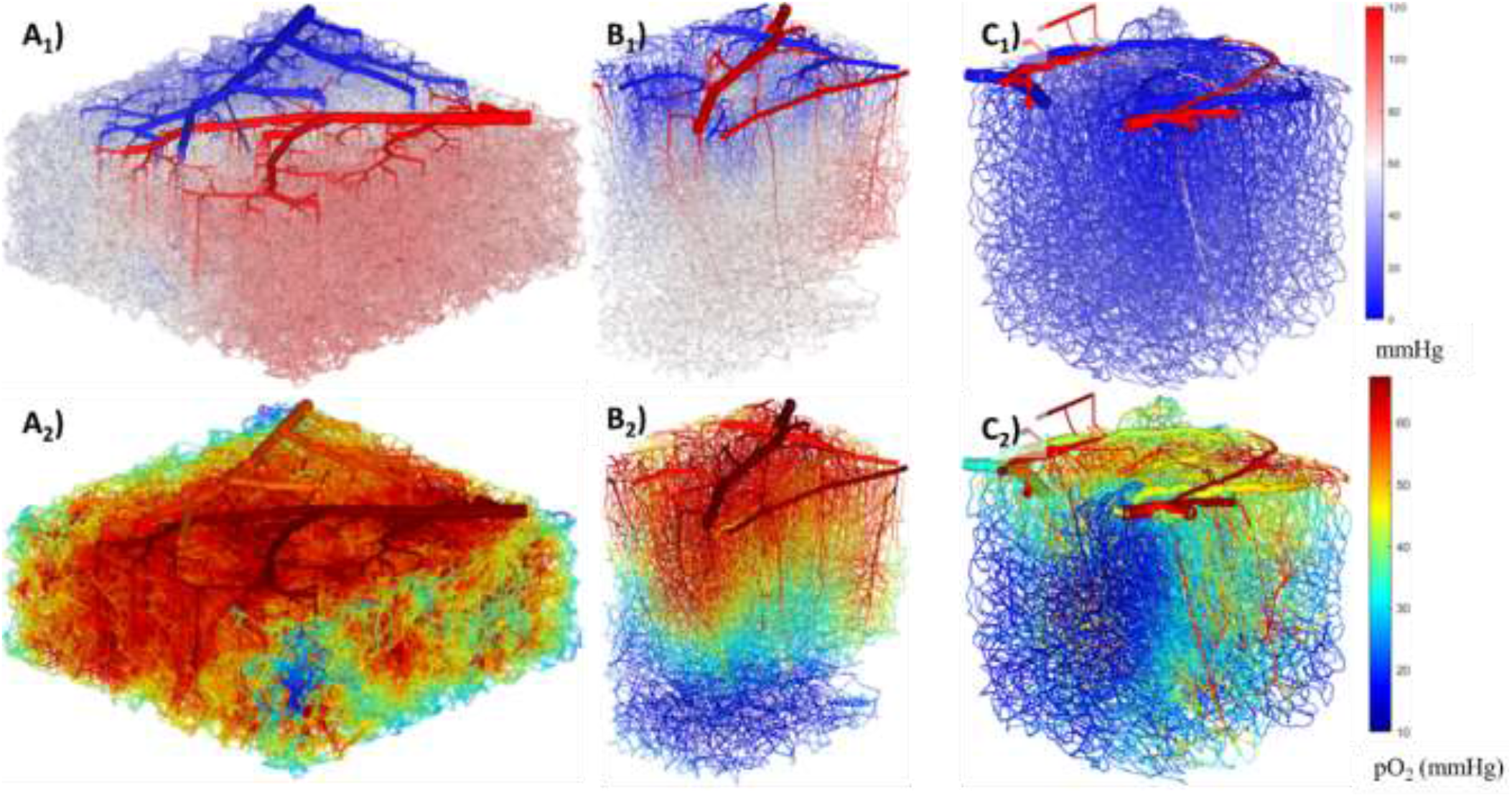
Voxelized synthetic and realistic in-vivo networks. Top-row: blood pressure in mmHg. Bottom row: steady state blood oxygen partial pressure in mmHg. A) 3 × 3 × 1.3 mm^3^ synthetic network [1] with 636,350 vessel segments. OEF = 12.3%. CMRO_2_ = 9.8e-06 mol/L/s B) 1.1 × 1.5 × 1.8 mm^3^ in-vivo network [23] with 39,547 vessel segments. OEF = 10.1%, CMRO_2_ = 3.2e-06 mol/L/s C) 1.1 × 1 × 1.1 mm^3^ in-vivo network [23] with 134,229 vessel segments. OEF = 34.6%, CMRO_2_ = 9.5e-06 mol/L/s.

With a solution **p** for the node pressures, we compute the blood flow in each edge from equation (3). At this point it may be advantageous to relabel the nodes of the vascular network such that the direction of each edge matches the direction of flow. Although this step is not strictly necessary, it has the advantage of turning the vascular graph into a directed acyclic graph (DAG) for which we can find a topological ordering of the nodes and for which the numerical flow values *q* are always positive.

### 2.4 O_2_ convection in the vascular domain

We assume a constant O_2_ concentration supplied to the inlet (=high pressure) nodes. The flow of O_2_ into a vascular control volume is given by the bulk blood inflow, and the flow out is given by the bulk blood outflow and by O_2_ diffusion through the vessel walls into the surrounding tissue. The latter is realized by the edges comprising the bipartite coupling graph, whereby every edge connecting a node to the tissue domain is an edge through which O_2_ can be transferred between domains, and is driven by the concentration gradient according to Fick’s law; the former is accomplished by blood flow through the vascular edges as established in section 2.3. The steady-state vascular concentrations are thus a balance between bulk flow in the vascular graph and diffusion through the vessel walls into the tissue graph: if mass transfer through the walls was not present, the O_2_ concentration would be constant throughout the vascular domain as the constant inlet value permeates through the whole network.

In the remainder of the paper, we will let **c**_*v*_ denote the length *n_p_* vector of steady state O_2_ concentrations in each nodal control volume and **c***_t_* denote the length *n_t_* vector of steady state concentrations of dissolved O_2_ in each tissue voxel. The equation for mass balance at steady state for all nodes in the network is

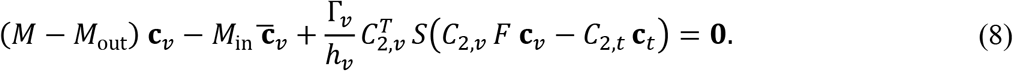

The first two terms of (8) express the flow balance in each nodal control volume due to bulk-flow (convection). The convection matrices *M, M*_out_, and *M*_in_ act on a vascular signal by returning the convective mass-balances in the interior, inlet, and outlet nodes respectively. The vector 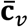 contains the values of the constant feed at the inlets.

The third term in (8) expresses the mass flux into tissue due to the local concentration difference between domains. The parameters Γ_*v*_ and *h_v_* are respectively the vessel wall permeability (in units of length^2^/time) and thickness. The diagonal matrix *S* contains the fractional surface area of each nodal control volume through which mass can diffuse. The matrices *C*_2,*v*_ and *C*_2,*t*_ act on the vascular and tissue concentration vectors respectively by projecting their values onto the surfaces of the nodal control volumes. Their transposes 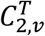 and 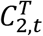 have the opposite effect, of projecting the surface values back onto the nodal control volumes and tissue voxels respectively. The diagonal operator *F* identifies the form of vascular O_2_ which can cross the vessel wall: in section 2.6 we suppose that O_2_ bound to hemoglobin can freely pass into the tissue, in which case *F* is the identity operator, whereas in section 2.7 we require O_2_ to be free in the plasma and model the dissociation kinetics with the Hill equation [18]. In the latter case *F* is a nonlinear operator.

We abbreviate the fully coupled convection equation in terms of the unknowns **c***_v_* and **c***_t_* as

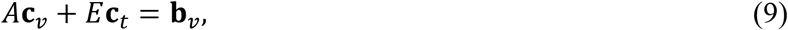

where

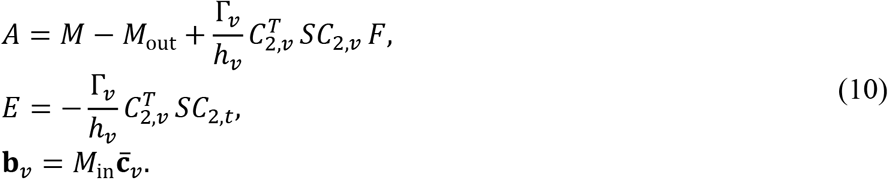

Note that *A* is a matrix (linear operator) if we ignore dissociation kinetics and a nonlinear operator otherwise. More detail on the construction of these matrices is presented in supplemental file S2.

### 2.5 O_2_ metabolism and diffusion in the tissue domain

Just as with the vascular domain, our equations ensure mass balance per unit time at steady state for each tissue voxel. Mass transfer in the tissue domain is achieved by diffusion between adjacent voxels, diffusion through the vascular walls via the edges of the bipartite coupling graph, and metabolic reactions.

The mass balance equation at steady state for all voxels is

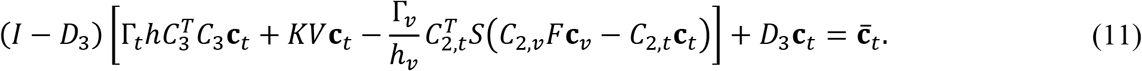

Since the decision matrix *D*_3_ identifies the boundary of the rectangular tissue domain and 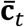 is a vector of prescribed boundary values, the first (bracketed) and second terms of (11) express net mass flux in the interior voxels and Dirichlet boundary conditions in the boundary voxels respectively.

The bracketed term of (11) contains three terms. The first term is a finite-volume discretization of the continuous Laplacian on a rectangular Cartesian grid: 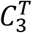 is the incidence matrix of the grid-graph, and so 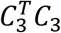 is its Laplacian matrix [31]. With **Γ**_*t*_ the diffusivity of O_2_ in tissue and *h* the constant voxel side-length, it follows that this term models diffusive flux of O_2_ through the voxel sides [33].

The second term in the brackets expresses consumption of O_2_ in tissue from metabolic reactions. We model reactions to first order, whereby the extinction rate per unit volume is directly proportional to the concentration. The diagonal matrices *K* and *V* contain the first order reaction rates (in units of 1/time) and the volumes of reacting tissue per voxel respectively. In our experiments we assume that the reaction rates have a constant positive value for all voxels and that the whole voxel is reactive, leading to a constant reacting volume of *h*^3^ per voxel. One possible alternative is to let the reacting volumes be the volume of the voxel minus the total volume of vascular control volumes intersecting with it, as in [11].

The third term in the brackets expresses the mass flux from the vascular domain: it closely mirrors the corresponding term in equation (8).

As in equation (9) we abbreviate the fully coupled reaction-diffusion equation (11) in terms of the unknowns **c***_v_* and **c**_*t*_ as

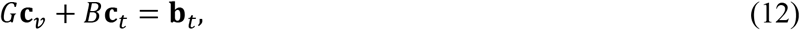

where

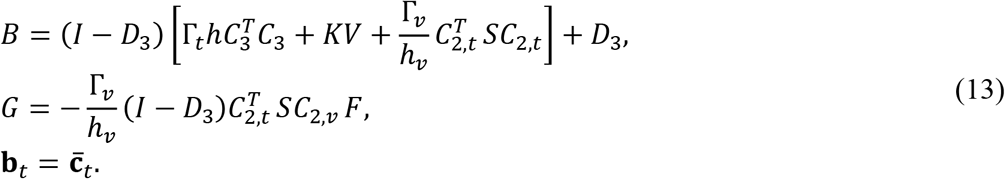

Note that *G* is a matrix (linear operator) if we ignore dissociation kinetics, and a nonlinear operator otherwise. More detail on the construction of these matrices is presented in supplemental file S2.

### 2.6 Blood-tissue coupling by domain decomposition with the Schur complement

#### 2.6.1 Poisson reactive diffusion with implicit vascular source terms

Previously in [1], systems (9) and (12) were solved simultaneously with an iterative algebraic solver. Here, we take advantage of the structure of the convection and reaction-diffusion matrices to build an even more efficient solution strategy.

As stated in the overview, the Poisson problem does not admit coupling of the diffusive-reactive tissue model to mass transfer from a vascular network, where oxygen is convected (=hyperbolic transport phenomenon). To preserve the desired structure for the elliptic tissue diffusion with reactive sink terms, we propose to incorporate oxygen supply from blood to tissue by introducing oxygen source terms. We use the Schur complement method [19] to obtain implicit expressions for unknown blood oxygen by performing partial Gauss elimination on the convective vascular oxygen equation subsystem. The Schur elimination of the vascular O_2_ tension can be interpreted as a separate solution for oxygen convection over a vascular network and is thus effectively a domain decomposition technique. Oxygen convection in blood can be effectively solved using graph matrix operations for finite volume discretization with upwinding [30, 33].

The resulting reduced ***Schur complement system*** can be solved the desired tissue O_2_. Once the tissue states are calculated, we recover vascular states from convection. We note that the overall master oxygen exchange problem is dominated by reactive diffusion in tissue as tissue cells which interact with vascular voxels are relatively sparse within the entire ensemble of tissue voxels, especially at high resolution. Thus, we propose to effectively precondition the Schur complement system with the solution to a pure diffusion problem with known source terms (a Poisson problem) as an approximation to unknown tissue states. Since the canonical Poisson problem is amenable to fast Fourier algorithms, the preconditioning step can be performed quickly, even for very large domains exceeding one billion equations without the need for supercomputers.

In section 2.6.2 we assume that O_2_ bound to hemoglobin can freely pass through the vessel wall into the extravascular tissue; consequently, the operator *F* in equations (10) and (13) is the identity operator, and the equations (9) and (12) are linear. Section 2.7 deals with nonlinear O_2_ dissociation kinetics which we solve with Newton’s method. The linear subproblems of section 2.6 are then applied within each Newton step.

#### 2.6.2 Implicit evaluation of O_2_ convection in blood

The master equation for the whole model is the pair of simultaneous equations (9) and (12), represented below as a block matrix equation:

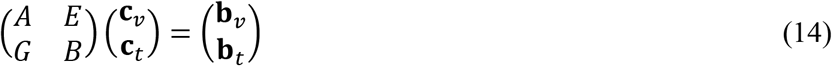

Solving for **c***_v_* in equation (9) we get

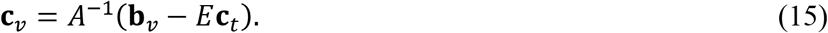

This expression for the vascular states implies that (12) can be reframed as a linear system for only tissue concentration **c***_t_* called the Schur complement system [19]:

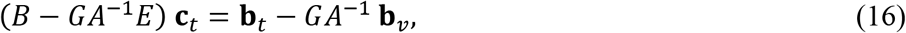

or

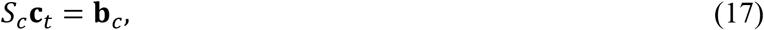

where

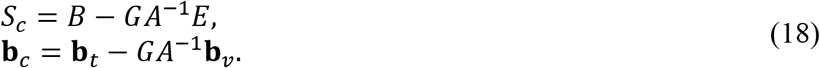

Note that the reduced system in equation (17) implicitly accounts for vascular state dependent mass transfer. The reduced system with Schur complement is equivalent to the fully coupled system in (12), without the need to iterate between unknown vascular and tissue states to compute oxygen exchange. Given a solution **c***_t_* to equation (17) we recover the corresponding vascular concentrations **c***_v_* from (15).

For the vascular state elimination strategy to be effective, the matrix *A* needs to be easily invertible. From its definition (10) note that *A* expresses convection through the vascular network with mass transfer into tissue, for which there always exists a unique solution. Moreover, the existence of a blood-flow solution in (7) implies that the flow network is a directed acyclic graph (DAG) when orienting edges of the vascular graph in the direction of blood flow. We should therefore expect to solve for the convected O_2_ states by mere forward propagation from the inlet nodes. Indeed, the nodes of a DAG can be topologically ordered: if *i* and *j* are nodes, then *i < j* if and only if there is a directed path from *i* to *j* [34]. It follows that under this re-labeling the matrix *A* is triangular, so the system of equations *A***u = v** can be solved for **u** cheaply by back-substitution: there is no need to compute the inverse matrix *A*^−1^ explicitly. See supplemental file S2 for discussion of the structure of the matrix *A:* the convection term is triangular under topological ordering whereas the mass-transfer term is diagonal.

Since computation of matrix by vector products *A*^−1^**v** by back-substitution is cheap, we implement the Schur complement matrix as a linear operator: from its definition (16) it follows that *S_c_* acts upon the tissue state by inducing a vascular state that is in convective and mass transfer balance with it and then superimposing the effect of diffusion. In this way the Schur complement implicitly supplements the diffusion dominated operator *B* with mass transfer from the vascular network which is consistent with the blood flow.

Although the system (17) has fewer unknowns than the full system (14), the Schur complement matrix *S_c_* is typically dense and extremely large: the number of tissue voxels is large compared to the number of vascular segments (up to 10^3^ times larger in our experiments) and so the time and memory complexity of the simulation is dominated by the number of tissue voxels, and hence the resolution of the voxel array. This necessitates a preconditioning step.

#### 2.6.3 Preconditioning the Schur complement system

We need to solve the linear Schur complement system (17) to obtain the tissue concentration. Iterative Krylov subspace methods require preconditioning to achieve rapid convergence. Following [20] we claim that if *P* is a good preconditioner for *B*, then *P* will be a good preconditioner for *S_c_* as well. The preconditioned system is

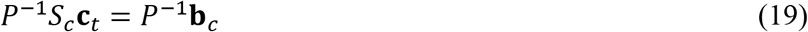

where *P* is a good preconditioner for *B*.

One way to obtain a good preconditioner is to approximate the matrix with another matrix which is easier to invert. The matrix *B* is essentially the matrix for a standard finite volume discretization of the continuous Poisson equation −∇^2^*u* = *ϕ* (i.e., a steady-state diffusion problem with constant source terms) with Dirichlet boundary conditions and added elements along the diagonal (it’s not hard to see that the matrix 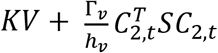 in (13) is diagonal: *KV* is diagonal by definition and the mass transfer between domains for a particular voxel does not depend on the values of any other tissue voxel). Hence, as suggested in [21] we take the preconditioner matrix to be the discretization operator for the Poisson equation with Dirichlet boundary conditions:

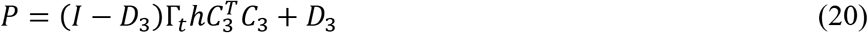

It follows that at every step of a Krylov subspace iteration finding the solution of (19) we must compute the matrix by vector product **u =** *P*^−1^**v** for an arbitrary vector **v** which, by definition, approximately solves the discrete Poisson equation with Dirichlet boundary conditions on a 3D Cartesian domain *P***u = v** [19].

For this to be an effective preconditioner we must be able to compute *P*^−1^**v** (solve a pure steady-state diffusion problem) quickly and efficiently for very large Cartesian domains. Fortunately, this can be done with *fast* discrete Fourier methods.

#### 2.6.4 Fast Poisson Algorithm

The pure steady-state diffusion problem is the continuous Poisson equation 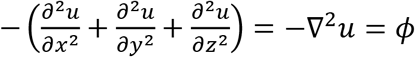. Suppose that its finite-volume discretization with Dirichlet boundary conditions on a 3D rectangular domain yields the matrix equation

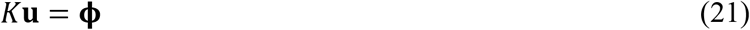

for a known scalar field **ϕ** and unknown vector of interior points **u**. Then, *K* has an eigendecomposition *K* = *W*Λ*W*, where *W*^−1^ = *W* and multiplication of an arbitrary vector by the modal matrix *W* is equivalent to taking the discrete sine transform (DST-I) of that vector [15, 16]. Consequently, a direct solution **u** to (21) can be computed as

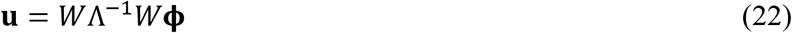

We can use the open-source library FFTW [16] to compute *W***b**, the DST-I of an arbitrary vector **b**, in *O*(*n* log *n*) operations; hence a fast solution (22) to the Poisson equation is obtained by taking a discrete sine transform of a 3D array, dividing by the known eigenvalues, and then taking another discrete sine transform. See supplemental file S3 for more details on the fast Poisson solver.

When solving the preconditioned Schur complement system (19) with a Krylov subspace method, at every iteration we compute the matrix by vector product, *P*^−1^**b**, for an arbitrary vector **b** by computing the corresponding solution to a discrete Poisson equation with Dirichlet boundary conditions with the fast Sine transform method outlined above. Together with the operator implementation of the Schur complement *S_c_* we can perform preconditioned Krylov subspace iterations for extremely large problem domains speedily and with a judicious utilization of memory. We show in section 3 that this is an effective preconditioner in the sense that the Krylov subspace iterative solver GMRES applied to the system (19) for the tissue oxygen values converges in much fewer iterations than it would when applied to the non-preconditioned system (17).

### 2.7 Nonlinear O_2_ dissociation kinetics

More realistic than the linear mass transfer model for oxygen exchange is the nonlinear dissociation kinetics [14]. Next, we consider convection of bound oxygen in the vascular domain but require that it dissociates from hemoglobin before it becomes available for diffusive transport into tissue. In particular, we suppose that the operator *F* in the trans-membrane mass transfer terms of (10) and (13) applies the nonlinear function *f* to every component of its input vector. The function *f* takes as input the concentration of oxygen bound to hemoglobin and outputs the corresponding concentration of oxygen dissolved in plasma under the assumption that oxygen is in equilibrium between its free and bound states. To model this relationship, we use the empirical Hill equation [18, 7]:

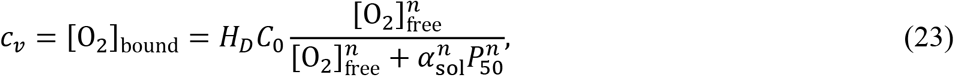

where *H_D_* is the discharge hematocrit (assumed constant throughout the network), *C*_0_ is the concentration of hemoglobin-bound oxygen in a fully saturated red blood cell, *n* is the empirical Hill coefficient, *P*_50_ is the pO_2_ at 50% saturation, and *α*_sol_ is the solubility of oxygen in plasma such that [O_2_]_free_ = *α*_sol_*P* for partial pressure *P*. This relation can be analytically inverted to yield the nonlinear relationship

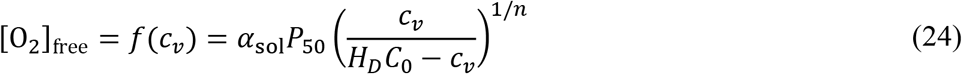

Since *f* is nonlinear it follows that the operator *F* and thus the master equation (14) is nonlinear. Therefore, we apply Newton’s method to attempt to find a solution [22]. Given an initial guess 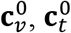, Newton’s method produces the sequences 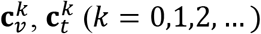 defined by

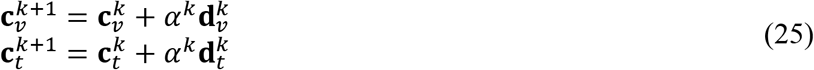

where the descent vectors 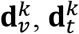 comprise a solution to the linear equation obtained by setting the first order Taylor expansion of the residual error of (14) equal to zero:

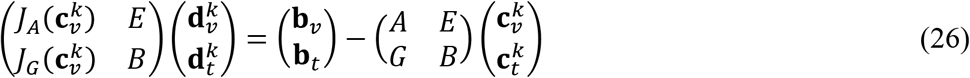

and the step length *α^k^* is chosen to approximately minimize the residual error of (14) on the line 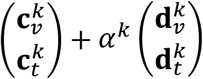. The Jacobian matrices *J_A_* and *J_G_* are given by

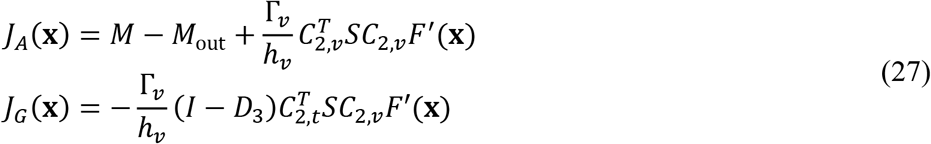

where the diagonal matrix 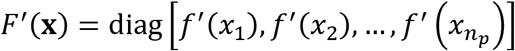 includes the derivative of the inverse Hill equation (24).

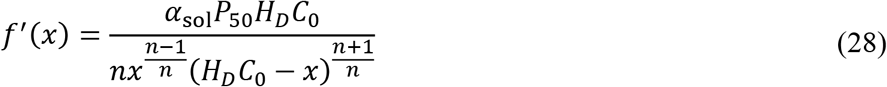

Since the matrix *B* in (26) is unchanged from section 2.6 we may construct a Schur complement system and use an identical preconditioner. In this manner we use the method of section 2.6 to solve (26) and find a downhill descent in Newton direction with *line search* in each iteration. To find suitable scalar acceleration *α^k^* we take the argmin of a quadratic approximation to the one-dimensional function

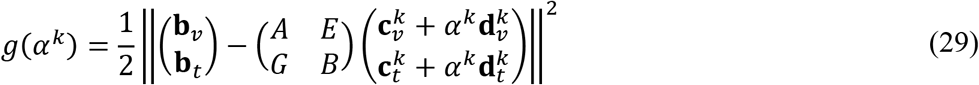

along the lines of Appendix C in [22]. Since the exponents involving the Hill coefficient *n* in (24) and (28) are not integers, the values of *c_v_* must be strictly constrained to the open interval (0, *H_D_C*_0_) to produce real-valued (and physiologically consistent) concentration fields. Depending on the initial guess, Newton’s method may produce components of **c***_v_* outside this range, so we simply map these values back into the feasible range before starting a new Newton iteration. In particular, for *i* ∈ {1,2,…, *n_p_*} if *c_v,i_* < 0 then 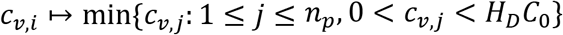 and if *c_v,i_* > *H_D_C*_0_ then 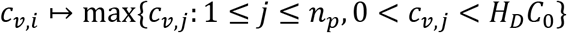.

## 3 Application of massive O_2_ exchange simulation in the cerebral cortex

All numerical experiments were performed in MATLAB version R2022a on an office PC with an Intel Xeon W-2125 processor at 4.01GHz and 256 GB of memory. For the fast-Poisson preconditioning we use the open-source FFTW library [16] to compute the discrete sine transform (DST-I) by way of a Matlab frontend [35]. By way of comparison, we note that it was found in [1] that for the 3×3×1.3 mm^3^ network in Figure 3A and about 95 million tissue voxels with linear mass flux, the automatic solver [36] converged in about 19 hours. With our improved methods, in this work we solved for 535 million tissue voxels with nonlinear mass flux on the same network and the same computer in one hour.

### 3.1 Pure diffusion of an ideal solute

To demonstrate the speed of the preconditioner described in section 2.6 we solve the finite difference discretization of the 3D Poisson equation (21) on an *n_x_* × *n_y_* × *n_z_* voxel rectangular domain with Dirichlet boundary conditions of zero and with a randomized right-side vector **ϕ**. For all cases under one billion voxels, we set *n_x_* = *n_y_* = 2^*k*^ – 1, and *n_z_* = 2^*k*–1^ – 1 for *k* = 5 through 10. Side lengths of this form are optimal for the fast transform algorithm. Solution times are compared in Table 1 and Figure 4 to different standard solvers. The results show that our method drives residuals to arbitrarily small values and can solve the Poisson problem in 452 seconds for a mesh with 1.5 billion mesh cells. Standard methods (mldivide, \ operation in Matlab) and PCG (conjugate gradient with modified incomplete Cholesky preconditioning) fail for domain sizes larger than 1 million equations.

**Table 1.** Performance metrics for the pure diffusion problem. The relative residual of (21) is defined as ||**ϕ** – *K***u**||/||**ϕ**||. “mldivide” is the standard matrix equation solver in Matlab, and PCG is the preconditioned conjugate gradient method. The preconditioning method used in PCG is the modified incomplete Cholesky factorization with no fill. An asterisk indicates that the solver did not converge in a reasonable time.

| num voxels (millions) | solution time (seconds) |  |  | relative residual |  |  |
| --- | --- | --- | --- | --- | --- | --- |
|  | fast | mldivide | PCG | fast | mldivide | PCG |
| 0.01 | 0.01 | 0.06 | 0.08 | 1e-14 | 2e-14 | 1e-10 |
| 0.1 | 0.02 | 1.4 | 0.3 | 5e-14 | 7e-14 | 8e-11 |
| 1 | 0.07 | 84 | 4 | 2e-13 | 3e-13 | 8e-11 |
| 8.3 | 0.4 | * | 48 | 9e-13 | * | 9e-11 |
| 66.6 | 3 | * | * | 4e-12 | * | * |
| 535 | 24 | * | * | 2e-11 | * | * |
| 1,002 | 126 | * | * | 4e-11 | * | * |
| 1,499 | 452 | * | * | 4e-11 | * | * |

**Figure 4.**
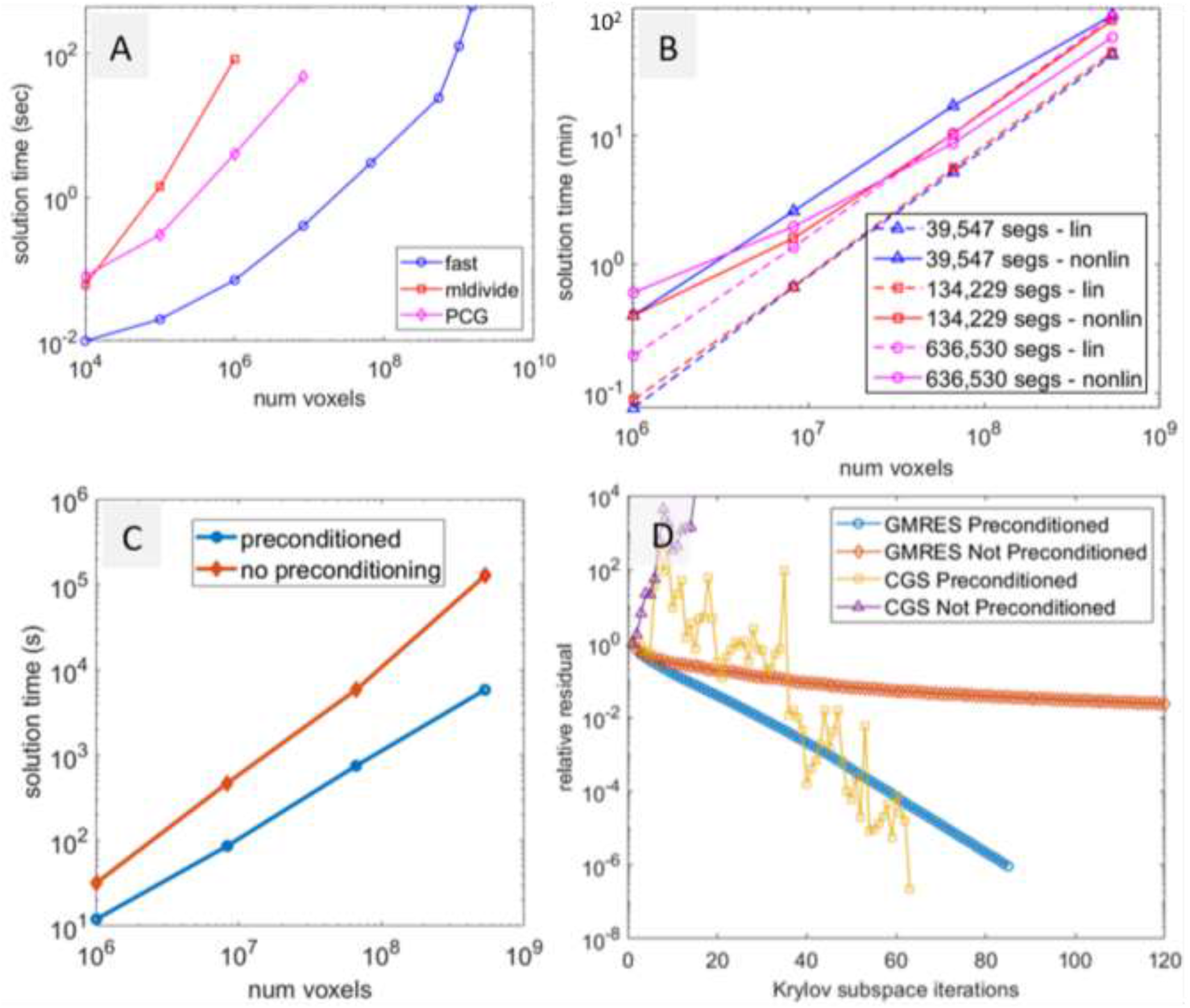
Comparison of solution times of the fast Poisson solver to a standard direct solver “mldivide” in Matlab and the iterative solver PCG (conjugate gradient with modified incomplete Cholesky preconditioning) for symmetric matrix systems. B) Comparison of solution times between the linear and nonlinear formulations of the O_2_ exchange for the three networks in Figure 2. C) Comparison of solution times for the linear problem with and without fast-Poisson preconditioning. D) The fast-Poisson preconditioning has a nontrivial effect on the size of the relative residual of the linear problem at each iteration, for two different Krylov subspace methods for square non-symmetric systems: GMRES (generalized minimal residual) and CGS (conjugate gradient squared).

### 3.2 Idealized exchange of O_2_ across the BBB (linear mass transfer)

We solve the preconditioned Schur complement system (19) associated with the networks in Figure 3. This corresponds to the case where hemoglobin bound O_2_ is free to cross into extravascular space. The mass transfer term between vascular and tissue domains and the corresponding master equation (14) is linear.

We use GMRES(10) [i.e. GMRES with restart every 10 iterations] for varying resolutions (values of *n_t_* and *h*) and the parameter values listed in Table 2. The highest resolution we were able to achieve was just over one billion tissue voxels because of limited memory in our standard desk PCs (Intel Xeon W-2125 processor at 4.01GHz and 256 GB of memory). We stop the iterative solver when the relative residual drops below 1e-6. In all cases the relative residual of the whole system (14) was orders of magnitude smaller than this, so presumably many fewer iterations are needed in practice. The time to solve for **c***_v_* once the solution **c***_t_* was in hand was negligible by comparison (less than one minute). Performance metrics for identical resolutions as those in section 3.1 are reported in Table 3. A comparison of solution times is displayed in Figure 4. We observe that faster convergence times are possible with a different Krylov subspace method; for instance, conjugate gradient squared (CGS) but decided to stick with GMRES because it was the most reliable, in contrast to CGS and other solvers which do not necessarily reduce the residual at each iteration.

**Table 2.** Parameter values used in simulation. Reaction rate was tweaked to obtain a minimum pO_2_ of no less than 10 mmHg in each network, in agreement with the range in [1].

| Parameter | Value | Units | Description | Reference |
| --- | --- | --- | --- | --- |
| $\mu$ | 3.2e-05 | mmHg·s | Dynamic viscosity of blood plasma | [37] |
| $C_0$ | 0.02 | Mol/L | Bound $O_2$ concentration of fully saturated RBCs | [38] |
| $H_D$ | 0.4 | | Discharge hematocrit | [38] |
| $n$ | 2.59 | | Hill coefficient | [38] |
| $P_{50}$ | 40.2 | mmHg | Dissolved $O_2$ partial pressure at 50% saturation | [38] |
| $\alpha_{sol}$ | 1.2e-06 | Mol/L/mmHg | Solubility of $O_2$ in plasma | [38] |
| $\bar{c}_v$ | 68 | mmHg | Arterial $pO_2$ (constant for all inlets, converted to $[O_2]_{free}$ by $\alpha_{sol}$ in the linear case and $[O_2]_{bound}$ by the Hill equation (23) in the nonlinear case) | [1] |
| $\bar{c}_t$ | 40 | mmHg | Ambient tissue $pO_2$ (constant Dirichlet boundary conditions, converted to concentration by $\alpha_{sol}$) | [14] |
| $\Gamma_v$ | 1.77e+03 | $\mu m^2/s$ | Permeability of vessel wall to $O_2$ | [1] |
| $h_v$ | 1 | $\mu m$ | Vessel wall thickness | [1] |
| $\Gamma_t$ | 1.8e+03 | $\mu m^2/s$ | Diffusivity of $O_2$ in tissue | [4] |
| $K$ | 0.4, 0.07, 0.25 | 1/s | First order reaction rate (for networks $A_2$, $B_2$ and $C_2$ in Figure 3 respectively) | |

**Table 3.** Performance metrics for the preconditioned Schur system (19) in the linear case and Newton’s method in the nonlinear case. Three different networks were placed in rectangular tissue domains with an equal number of voxels. For *k* = 7,8,9,10, the number of voxels parallel to the *x* and *y* axes are 2*^k^* – 1, and the number of voxels parallel to the *z* axis is 2^*k*–1^ – 1 since the performance of the FFTW algorithm is optimized for side lengths of this form [16]. The linear Schur complement system and Newton subproblems were solved with GMRES(10) and stopped when the relative residual dropped below 1e-6. For the 1 billion voxel case we used GMRES(5) to circumvent memory constraints.

| Network size<br>(mm<sup>3</sup>) | Num<br>segments | Num voxels $n_t$<br>(millions) | Voxel side<br>length $h$ (μm) | Newton<br>iterations | Solution time -<br>Linear (min) | Solution time -<br>Nonlinear (min) |
| --- | --- | --- | --- | --- | --- | --- |
| 3 x 3 x 1.3 | 636,350 | 1 | 25 | 8 | 0.2 | 0.6 |
|  |  | 8.3 | 13 | 3 | 1.4 | 2 |
|  |  | 66.6 | 6.3 | 2 | 10.5 | 8.8 |
|  |  | 535 | 3.2 | 2 | 85.6 | 58.7 |
|  |  | **1,002** | **2** | **4** | **442 (7.4 hr)** | **435 (7.3 hr)** |
| 1.1 x 1 x 1.1 | 134,229 | 1 | 19 | 8 | 0.09 | 0.4 |
|  |  | 8.3 | 9.3 | 3 | 0.7 | 1.6 |
|  |  | 66.6 | 4.7 | 3 | 5.6 | 10.4 |
|  |  | 535 | 2.3 | 3 | 45 | 80.5 |
| 1.1 x 1.5 x 1.8 | 39,547 | 1 | 31 | 8 | 0.08 | 0.4 |
|  |  | 8.3 | 15 | 6 | 0.7 | 2.6 |
|  |  | 66.6 | 7.7 | 5 | 5.2 | 17.2 |
|  |  | 535 | 3.8 | 3 | 43 | 87.2 |

### 3.3 Cortical O_2_ extraction with nonlinear dissociation kinetics

In this section we use three networks derived from synthetic and in-vivo data whose voxelized representations are shown in Figure 3. The solution **c***_v_* is reflected in the coloration of the networks on the bottom row of this figure.

We apply Newton’s method to the full nonlinear system with resolutions identical to those in sections 3.1 and 3.2 and use preconditioned GMRES(10) at each Newton step to find the descent vector, as described in section 2.7. For the lowest resolution cases we use a constant vector for an initial guess; otherwise, we interpolate the solution from the closest lower resolution level to the higher resolution grid. For instance, the *n_t_* = 8.3 million case uses the interpolated output of the *n_t_* = 1 million case as its initial guess. Performance metrics are reported in Table 3. Solution time is cumulative, reflecting the time needed to compute the initial guess by solving the lower resolution cases. Planar cuts of the 3D solution in the representative *n_t_* = 66.6 million case are shown in Figure 5 and Figure 6, and a video showing the O_2_ field at all constant *z* values is shown in supplementary file S4. Figure 5 shows that features of the tissue O_2_ field are maintained between resolution levels, demonstrating mesh independence, and Figure 6 shows the level of detail achieved at even modest solution times, as the nonlinear 66.6 million voxel case converges in only 9 minutes. Physiological validation studies of our O_2_ model against two-photon imaging of the rat brain are presented in [1].

**Figure 5.**
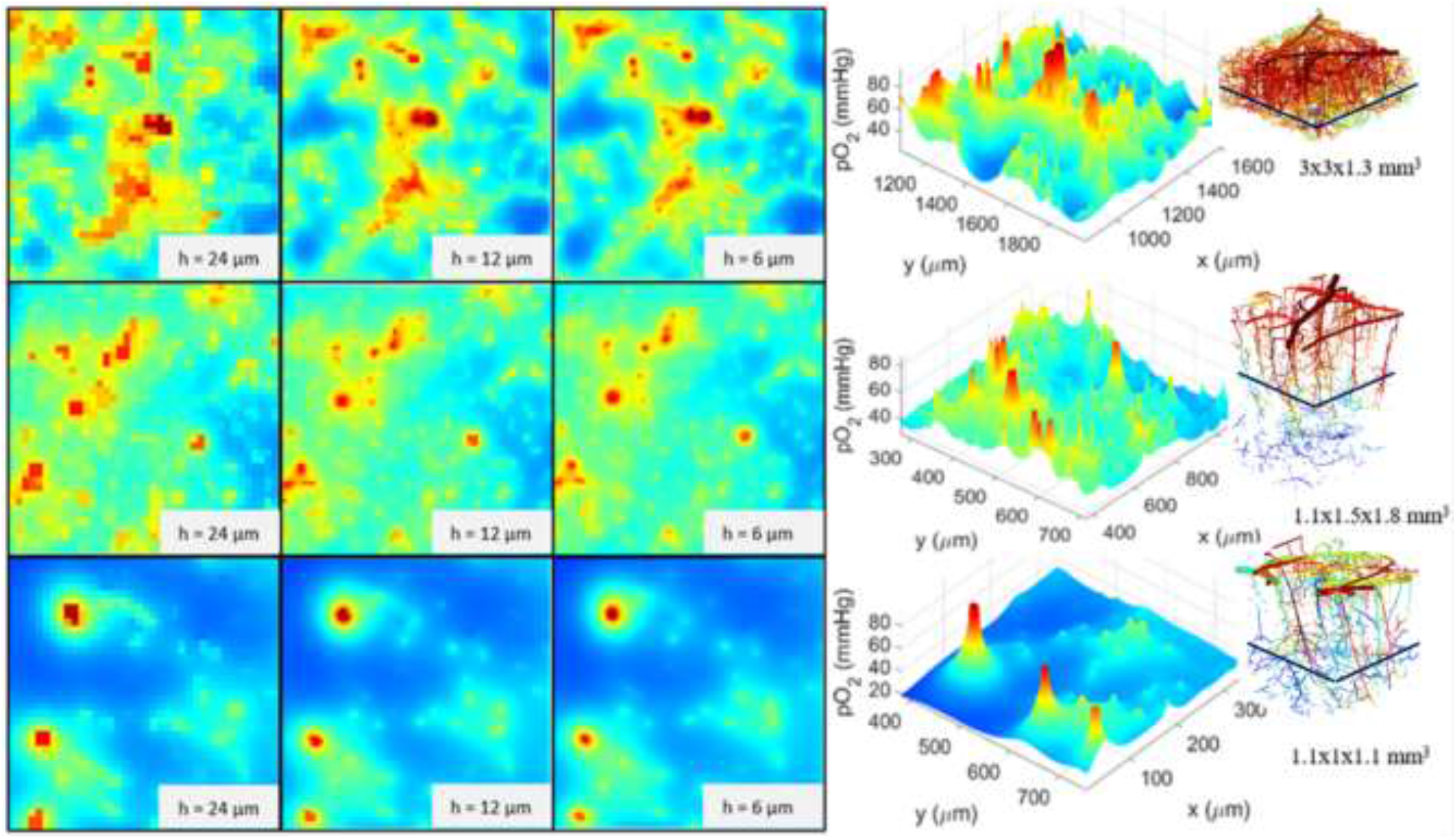
Comparison of features of the tissue pO_2_ field in a constant **z** slice at three different resolutions for three different networks. Features persist and become more sharply defined at higher resolutions, demonstrating mesh independence of the model. It is easy to see the characteristically steep and radially symmetric profile curves around penetrating arterioles when tissue pO_2_ is plotted as a surface.

**Figure 6.**
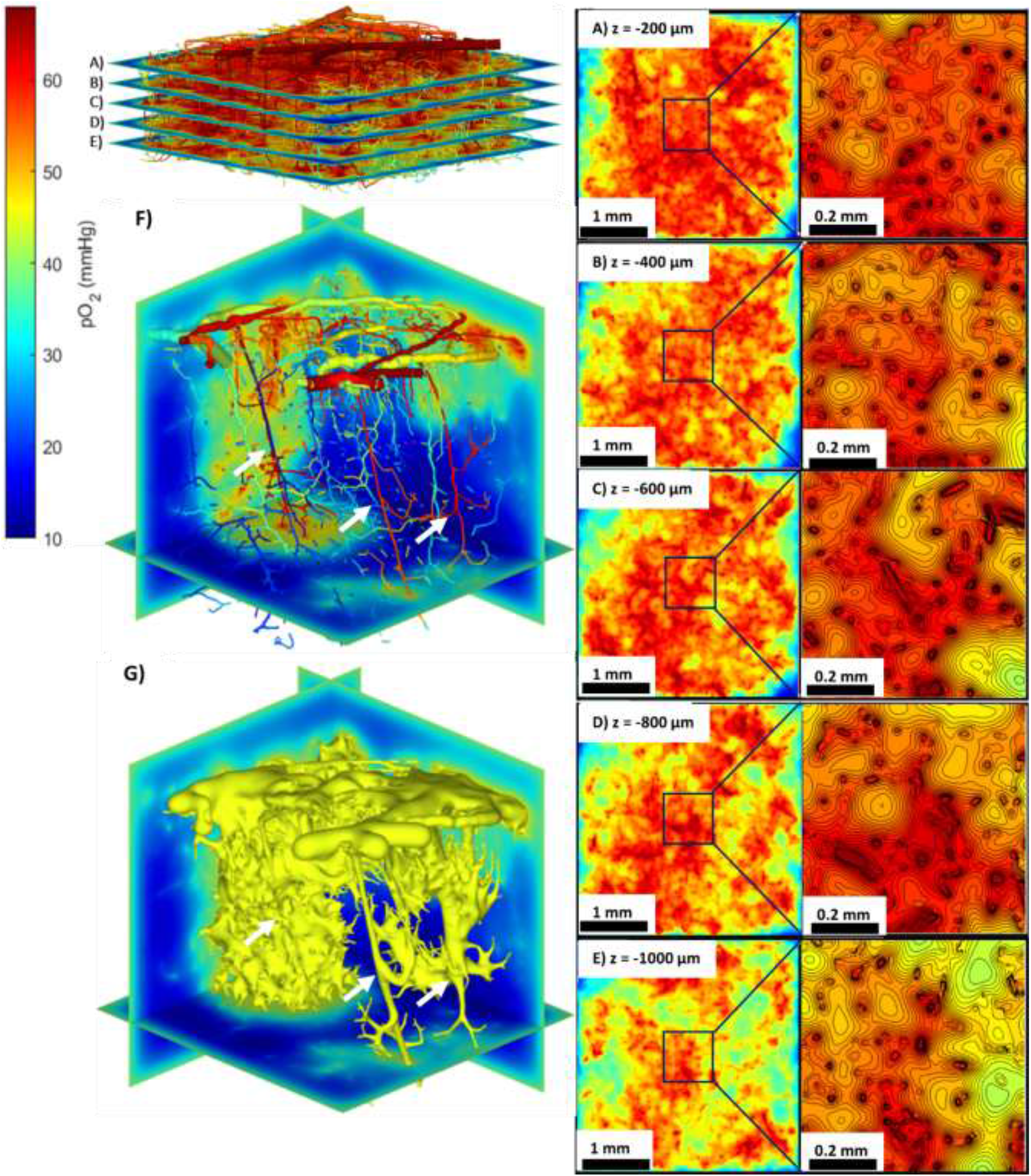
A-E) Horizontal cross-sections of O_2_ field about the 3×3×1.3 mm^3^ synthetic network with voxel side length of 6 μm. Zoom in on features demonstrates the level of detail acquired at modest resolution and solution times (about 10 minutes) for large networks. F) 3D rendering of the 1.1×1×1.1 mm^3^ mouse cortical network with vessels less than 5μm in diameter omitted and cross-sections of corresponding O_2_ field at 6 μm resolution. Penetrating arterioles are indicated with arrows. G) The same view as in F with the 45 mmHg isosurface of the O_2_ field displayed. The field clearly conforms to the shape of the network in 3D, especially near penetrating arterioles.

## 4 Conclusions

High resolution (up to the micron level) simulation of cortical oxygen perfusion is necessary for physiologically meaningful analysis of the relationship of microvascular topology to neurovascular dysfunction. We presented a novel mesh-free mixed-domain technique based on fast discrete sine transfer (DST) preconditioning which enabled the modeling of oxygen perfusion in cerebrovascular systems of unprecedented high resolution and anatomical complexity. This allows us to resolve the smallest scales of the microvasculature in realistic networks down to the single micron scale. Large scale domain sizes are also essential for eliminating non-physical boundary effects; this can be achieved by simulating oxygen exchange over a larger region but restricting analysis only to a small core region of interest far removed from the domain boundaries.

Our innovations rested on a dual mesh technique which uses graph representations of the vascular and tissue domains and their coupling for automatic generation of model equations. A Cartesian reference mask for the tissue domain with embedded vascular network enables a simple coupling scheme that can resolve the smallest capillaries without the need for fitted mesh generation. Furthermore, the Cartesian structured matrices allowed us to leverage fast Fourier transform algorithms to solve closely related Poisson problems extremely quickly at ultra-high resolution, affording a cheap preconditioner for the tissue oxygen computation. Finally, the Schur complement implicitly incorporates the effect of mass transfer from vasculature into tissue and generates a fully coupled system that can be effectively preconditioned by the fast Poisson solver. Nonlinear dissociation kinetics did not diminish the efficacy of the method, requiring only the additional complication of utilizing Newton’s method in which the linear subproblem is solved at each iteration.

The encouraging results for stationary conditions further motivate extension to the unsteady case for observing time dependent oxygen perfusion subject to stimuli such as neuronal firing or periodic excitation (e.g. vasomotor drive).

## Acknowledgments

The authors would like to gratefully acknowledge NIH for their financial support through projects 1R56AG066634-01 and 1R01AG079894-01 Models for aging and Alzheimer with image-based cerebrovascular network synthesis (iCNS), and 1U19NS123717-01.

## Appendix Table of Symbols

Appendix - Table of Symbols
| Symbol | Description |
| --- | --- |
| $n_p$ | Number of VAN nodes = control volumes |
| $n_f$ | Number of VAN edges = vessel segments |
| $n_t$ | Number of Cartesian grid points = tissue voxels |
| $n_c$ | Number of coupling edges = mass transfer edges between vascular and tissue domains |
| $C_1$ | $n_f \times n_p$ connectivity matrix for vascular network |
| $C_2$ | $n_c \times (n_p + n_t)$ connectivity matrix for bipartite coupling graph (all edges connecting vascular and tissue domains) |
| $C_{2,v}$ | First $n_p$ columns of $C_2$ |
| $C_{2,t}$ | Last $n_t$ columns of $-C_2$ |
| $C_3$ | Connectivity matrix for the grid graph of the tissue domain |
| $D_1$ | $n_p \times n_p$ decision matrix for the vascular boundary nodes |
| $D_3$ | $n_t \times n_t$ decision matrix for the tissue boundary nodes |
| $d_k$ | Diameters of VAN edges = vessel segments, $k = 1, 2, \dots, n_f$ |
| $L_k$ | Lengths of VAN edges = vessel segments, $k = 1, 2, \dots, n_f$ |
| $w_k$ | Reciprocal of hydraulic resistance in VAN edges (2), $k = 1, 2, \dots, n_f$ |
| $W$ | $n_f \times n_f$ diagonal matrix of $w_k$ values |
| $p_i$ | Blood pressure per vascular control volume, $i = 1, 2, \dots, n_p$ |
| $\mathbf{p}$ | $n_p \times 1$ vector of $p_i$ values (6) |
| $\bar{\mathbf{p}}$ | $n_p \times 1$ vector of boundary $p$ values (zero at interior nodes) |
| $q_k$ | Volumetric blood flow through VAN edges ($\mu\text{m}^3/\text{s}$), $k = 1, 2, \dots, n_f$ |
| $\mathbf{q}$ | $n_f \times 1$ vector of $q_k$ values (3) |
| $S$ | $n_c \times n_c$ diagonal matrix of $s_m$ values |
| $f$ | Real valued function mapping hemoglobin bound oxygen concentration to free oxygen concentration (24) |
| $F$ | Diagonal operator applying $f$ to every component of input vector |
| $M$ | Convection matrix |
| $M_{\text{in}}, M_{\text{out}}$ | Inlet and outlet convection matrices |
| $c_{v,i}$ | Average bound oxygen concentration in vascular nodal control volumes, $i = 1, 2, \dots, n_p$ |
| $\mathbf{c}_v$ | $n_p \times 1$ vector of $c_{v,i}$ values |
| $\bar{\mathbf{c}}_v$ | $n_p \times 1$ vector of prescribed constant vascular inlet concentrations (zero not at inlet nodes) |
| $K$ | $n_t \times n_t$ diagonal matrix of first order reaction rates (1/time) |
| $V$ | $n_t \times n_t$ diagonal matrix of reacting voxel sub volumes |
| $c_{t,r}$ | Average oxygen concentration in tissue voxels, $r = 1, 2, \dots, n_t$ |
| $\mathbf{c}_t$ | $n_t \times 1$ vector of $c_{t,r}$ values |
| $\bar{\mathbf{c}}_t$ | $n_t \times 1$ vector of prescribed constant tissue boundary values (zero at interior) |
| $A$ | Vessel on vessel dependence matrix (10) |
| $E$ | Vessel on tissue dependence matrix (10) |
| $\mathbf{b}_v$ | Right side of master vascular equation (10) |
| $B$ | Tissue on tissue dependence matrix (13) |
| $G$ | Tissue on vessel dependence matrix (13) |
| $\mathbf{b}_t$ | Right side of master tissue equation (13) |
| $S_c$ | Schur complement matrix (18) |
| $\mathbf{b}_c$ | Right side of Schur complement system (17), (18) |
| $P$ | Preconditioning matrix (20) |
| $\mu$ | Dynamic viscosity of blood plasma (mmHg.s) |
| $C_0$ | Bound $\text{O}_2$ concentration of fully saturated RBCs (Mol/L) |
| $H_D$ | Discharge hematocrit |
| $n$ | Hill coefficient |
| $P_{50}$ | Dissolved $\text{O}_2$ partial pressure at 50% saturation (mmHg) |
| $\alpha_{\text{sol}}$ | Solubility of $\text{O}_2$ in plasma (Mol/L/mmHg) |
| $\Gamma_v$ | Permeability of vessel wall to $\text{O}_2$ ($\mu\text{m}^2/\text{s}$) |
| $h_v$ | Vessel wall thickness ($\mu\text{m}$) |
| $\Gamma_t$ | Diffusivity of $\text{O}_2$ in tissue ($\mu\text{m}^2/\text{s}$) |
| $h$ | Voxel side length ($\mu\text{m}$) |

## References

[1] Hartung G, Badr S, Moeini M, et al. Voxelized simulation of cerebral oxygen perfusion elucidates hypoxia in aged mouse cortex. PLoS computational biology. 2021;17(1). doi:10.1371/journal.pcbi.1008584.

[2] Krogh A. The number and distribution of capillaries in muscles with calculations of the oxygen pressure head necessary for supplying the tissue. The Journal of Physiology. 1919;52(6):409–415. doi:10.1113/jphysiol.1919.sp001839.

[3] Reneau Jr DD, Bruley DF, Knisely MH. A digital simulation of transient oxygen transport in capillary-tissue systems (cerebral grey matter). Development of a numerical method for solution of transport equations describing coupled convection-diffusion systems. AlChE Journal. 1969;15(6):916–925. doi:10.1002/aic.690150620.

[4] Mintun MA, Lundstrom BN, Snyder AZ, Vlassenko AG, Shulman GL, Raichle ME. Blood flow and oxygen delivery to human brain during functional activity: Theoretical modeling and experimental data. Proceedings of the National Academy of Sciences. 2001;98(12):6859–6864. doi:10.1073/pnas.111164398.

[5] Buxton RB, Frank LR. A Model for the Coupling Between Cerebral Blood Flow and Oxygen Metabolism During Neural Stimulation. Journal of Cerebral Blood Flow and Metabolism. 1997;17(1):64–72. doi:10.1097/00004647-199701000-00009.

[6] Jespersen SN, Østergaard L. The roles of cerebral blood flow, capillary transit time heterogeneity, and oxygen tension in brain oxygenation and metabolism. Journal of Cerebral Blood Flow & Metabolism. 2012;32(2):264–277. doi:10.1038/jcbfm.2011.153.

[7] Secomb TW, Hsu R, Park EYH, Dewhirst MW. Green’s Function Methods for Analysis of Oxygen Delivery to Tissue by Microvascular Networks. Annals of Biomedical Engineering. 2004;32(11):1519–1529. doi:10.1114/B:ABME.0000049036.08817.44.

[8] Serajelahi B, Kharche S, Goldman D. Steady-State Tissue Oxygen Distributions Calculated by a Green’s Function Method and a Finite Difference Method: A Comparison. 42nd Annual International Conference of the IEEE Engineering in Medicine & Biology Society (EMBC). 2020:2279–2282. doi:10.1109/EMBC44109.2020.9175901.

[9] Celaya-Alcala JT, Lee GV, Smith AF, et al. Simulation of oxygen transport and estimation of tissue perfusion in extensive microvascular networks: Application to cerebral cortex. Journal of Cerebral Blood Flow and Metabolism. 2021;41(3):656–669. doi:10.1177/0271678X20927100.

[10] Fang Q, Sakadžić S, Ruvinskaya L, Devor A, Dale AM, Boas DA. Oxygen advection and diffusion in a threedimensional vascular anatomical network. Optics express. 2008;16(22):17530–17541. doi:10.1364/OE.16.017530.

[11] Linninger AA, Gould IG, Matinnan T, Hsu CY, Chojecki M, Alaraj A. Cerebral microcirculation and oxygen tension in the human secondary cortex. Annals of biomedical engineering. 2013;41(11):2264–2284. doi:10.1007/s10439-013-0828-0.

[12] Gould IG, Tsai P, Kleinfeld D, Linninger AA. The capillary bed offers the largest hemodynamic resistance to the cortical blood supply. Journal of Cerebral Blood Flow and Metabolism. 2017;37(1):52–68. doi:10.1177/0271678X16671146.

[13] Kuchta M, Laurino F, Mardal KA, Zunino P. Analysis and approximation of mixed-dimensional PDEs on 3D-1D domains coupled with Lagrange multipliers. SIAM Journal on Numerical Analysis. 2021;59(1):558–582. doi:10.1137/20M1329664

[14] Gould IG, Linninger AA. Hematocrit distribution and tissue oxygenation in large microcirculatory networks. Microcirculation. 2015;22(1):1–18. doi:10.1111/micc.12156.

[15] Golub GH, Van Loan CF. Matrix computations. 4^th^ ed. Baltimore, MD: The Johns Hopkins University Press; 2013.

[16] Frigo M, Johnson SG. The design and implementation of FFTW3. Proc. IEEE. 2005;93(2):216–231. doi:10.1109/JPROC.2004.840301.

[17] Briggs WL, Henson VE, McCormick SF. A multigrid tutorial. 2^nd^ ed. Philadelphia, PA: Society for Industrial and Applied Mathematics; 2000.

[18] Hill AV. The possible effects of the aggregation of the molecules of haemoglobin on its dissociation curves. The Journal of Physiology. 1910;40:4–7.

[19] Saad Y. Iterative methods for sparse linear systems. 2^nd^ ed. Philadelphia, PA: Society for Industrial and Applied Mathematics; 2003. doi:10.1137/1.9780898718003.

[20] Mandel J. On block diagonal and Schur complement preconditioning. Numerische Mathematik. 1990;58(1):79–93. doi:10.1007/BF01385611.

[21] Chan RH, Wong CK. Sine Transform Based Preconditioners for Elliptic Problems. Numerical Linear Algebra with Applications. 1997;4(5):351–368. doi:10.1002/(SICI)1099-1506(199709/10)4:5<351::AID-NLA103>3.0.CO;2-4.

[22] Bertsekas DP. Nonlinear Programming. 3^rd^ ed. Nashua, NH: Athena Scientific; 2016.

[23] Blinder P, Philbert TS, Kaufhold JP, Knutsen PM, Suhl H, Kleinfeld D. The cortical angiome: An interconnected vascular network with noncolumnar patterns of blood flow. Nature Neuroscience. 2013;16(7):889–897. doi:10.1038/nn.3426.

[24] Philbert TS, Kaufhold JP, Blinder P, et al. Correlations of Neuronal and Microvascular Densities in Murine Cortex Revealed by Direct Counting and Colocalization of Nuclei and Vessels. Journal of Neuroscience. 2009;29(46):14553–14570. doi:10.1523/JNEUROSCI.3287-09.2009

[25] Shih AY, Driscoll JD, Drew PJ, Nishimura N, Schaffer CB, Kleinfeld D. Two-Photon Microscopy as a Tool to Study Blood Flow and Neurovascular Coupling in the Rodent Brain. Journal of Cerebral Blood Flow & Metabolism. 2012;32(7):1277–1309. doi:10.1038/jcbfm.2011.196

[26] Kaufhold JP, Tsai PS, Blinder P, Kleinfeld D. Vectorization of optically sectioned brain microvasculature: Learning aids completion of vascular graphs by connecting gaps and deleting open-ended segments. Medical Image Analysis. 2012;16(6):1241–1258. doi:10.1016/j.media.2012.06.004

[27] Weisstein EW. Grid Graph. From MathWorld--A Wolfram Web Resource. https://mathworld.wolfram.com/GridGraph.html. Accessed June 13, 2022.

[28] Weisstein EW. Bipartite Graph. From MathWorld--A Wolfram Web Resource. https://mathworld.wolfram.com/BipartiteGraph.html. Accessed June 14, 2022.

[29] Weisstein EW. Incidence Matrix. From MathWorld--A Wolfram Web Resource. https://mathworld.wolfram.com/IncidenceMatrix.html. Accessed June 13, 2022.

[30] Park CS, Hartung G, Alaraj A, Du X, Charbel FT, Linninger AA. Quantification of blood flow patterns in the cerebral arterial circulation of individual (human) subjects. International journal for numerical methods in biomedical engineering. 2020;36(1):e3288. doi:10.1002/cnm.3288

[31] Chung FR. Spectral graph theory. Providence, RH: American Mathematical Society; 1997.

[32] Spielman DA. Spectral and Algebraic Graph Theory Incomplete Draft. 2019. http://cs-www.cs.yale.edu/homes/spielman/sagt/sagt.pdf. Accessed June 15, 2022.

[33] Patankar SV. Numerical heat transfer and fluid flow. Boca Raton, FL: CRC Press; 1980.

[34] Bang-Jensen J, Gutin GZ. Acyclic Digraphs. In: Digraphs: Theory, Algorithms and Applications. 2^nd^ ed. London: Springer-Verlag; 2009:32–34.

[35] Treeby B. MATLAB Discrete Trigonometric Transform Library. https://github.com/ucl-bug/matlab-dtts/releases/tag/v1.1. Accessed November 11, 2022.

[36] Balay S, Abhyankar S, Adams MF, et al. PETSc Web page. https://petsc.org/. Accessed December 31, 2022.

[37] Klabunde RE. Cardiovascular Physiology Concepts. https://cvphysiology.com/Hemodynamics/H011. Accessed July 17, 2022.

[38] Gagnon L, Sakadžić S, Lesage F, et al. Quantifying the Microvascular Origin of BOLD-fMRI from First Principles with Two-Photon Microscopy and an Oxygen-Sensitive Nanoprobe. The Journal of Neuroscience. 2015;35(8):3663–3675. doi:10.1523/JNEUROSCI.3555-14.2015.

